# Integrative analyses to investigate the link between microbial activity and metabolites degradation during anaerobic digestion

**DOI:** 10.1101/2020.02.13.946970

**Authors:** Laetitia Cardona, Kim Anh Lê Cao, Francesc Puig-Castellví, Chrystelle Bureau, Céline Madigou, Laurent Mazéas, Olivier Chapleur

## Abstract

Anaerobic digestion (AD) is a promising biological process which converts waste into sustainable energy. To fully exploit AD’s capability, we need to deepen our knowledge of the microbiota involved in this complex bioprocess. High-throughput methodologies open new perspectives to investigate AD process at the molecular level, supported by recent data integration methodologies to extract relevant information. In this study, we investigated the link between microbial activity and substrate degradation in a lab-scale anaerobic co-digestion experiment, where bioreactors were fed with 9 different mixtures of three co-substrates (fish waste, sewage sludge, and grass). Samples were profiled using 16S rRNA sequencing and untargeted metabolomics. In this article, we propose a suite of multivariate tools to statistically integrate these data and identify coordinated patterns between groups of microbial and metabolic profiles specific of each co-substrate. Five main groups of features were successfully evidenced, including cadaverine degradation found to be associated with the activity of microorganisms from the order *Clostridiales* and the genus *Methanosarcina*. This study highlights the potential of data integration towards a comprehensive understanding of AD microbiota.

## Introduction

Deciphering the microbial communities in diverse domains, such as health, food safety and environment, has been widely addressed by the development of high-throughput technologies. In particular, Anaerobic Digestion (AD) bioprocess is constituted by extremely complex microbial communities, mainly composed of bacteria and archaea, with high functional redundancies. While AD is a promising bioprocess for the production of biogas, a sustainable energy, from the degradation of organic waste, the key microorganisms driving AD remain largely unknown. AD bioprocess involves four principal steps: hydrolysis, acidogenesis, acetogenesis and methanogenesis. The optimal functioning of AD relies on the efficient network between the microbes carrying successively these steps. However, the microbial diversity and interactions between microorganisms are highly sensitive and depend on multiple technical parameters such as temperature (Madigou et al., 2019; Noll et al., 2010), presence of inhibitors (Li et al., 2017; Poirier et al., 2016; Sousa et al., 2013), or feedstock composition (Zamanzadeh et al., 2017).

Emerging omics and high-throughput approaches can monitor microbial complexity across multiple functional levels (Vanwonterghem et al., 2014). Metagenomics, metatranscriptomics, metaproteomics and metabolomics provide necessary information related to a community’s genes, gene expression, proteins and metabolite production. They reveal potential and activated functions of microbial communities. In our context, these omics can unravel the intricate networks of functional processes of AD, especially when comparing bioreactors under different operational conditions to elucidate their functioning (Carballa et al., 2015; De Vrieze and Verstraete, 2016; Lü et al., 2014).

In parallel to the important advancements in these fields, the recent development of specific statistical methods and user-friendly workflows has substantially improved the analysis and visualisation of these omics results (Bouhlel et al., 2018; Callahan et al., 2016; Rohart et al., 2017; Singh et al., 2019). As the statistical analysis of single omics is not sufficient to decipher complex microbial relationships, the combination of information from several data sources has become mandatory. Computational analytical methods have a rising potential to capitalise on this rich data but they are still at their infancy and are not broadly used for this type of problems. Classical multivariate methods, including Principal Component Analysis (PCA) or Principal Coordinate Analysis (PCoA), have been widely used for single omics datasets, such as 16S or metabolomic data (Poirier et al., 2016; Wang et al., 2017). However, such methods are exploratory and do not fully characterised the microbial communities playing a role in the biological systems being studied. Moreover, these methods are not currently used to integrate information from different omics experiments. Novel computational and statistical methods are becoming available to fully harvest the large amount of data generated in multi-omics experiments (Lê Cao et al., 2009; Singh et al., 2019; Straube et al., 2017). These methods aim to extract complementary information from several datasets for a holistic understanding of the interplay between the different levels of information.

Several AD studies have already shown the utility of using omics technologies to decipher the anaerobic microbial population (Amha et al., 2018; Hassa et al., 2018). Analyses of a single omics are routinely carried out (Bize et al., 2015; Cai et al., 2016) but the use of several paired omics on the same samples, and their statistical integration are rare currently. For example, Beale *et al.* applied metagenomic and metabolomic approaches to obtain new insights on the diversity and activity of the anaerobic population after stress (Beale et al., 2016) but no data integration was conducted between the two approaches. However, correlation between transcriptomics and metabolomics has already been used to identify important metabolites and genes for example in fungal or plants (Askenazi et al., 2003; Hoefgen and Nikiforova, 2008).

Our study is the first of its kind in the anaerobic bioprocess field to investigate the link between microbial activity from 16S rRNA sequencing data and patterns of substrate degradation from metabolomics data. We propose a statistical framework to integrate these datasets to identify associations, or correlations, between microbial activity and metabolites specific of substrates degradation. To that end, 27 anaerobic bioreactors were fed with binary mixtures of three substrates of different chemical composition (sludge and grass, or sludge and fish) at different proportions. The microbial community was analysed through the sequencing of the 16S rRNA to characterise metabolically active microorganisms (De Vrieze et al., 2018). The patterns of the substrate degradation were studied with untargeted metabolomics using LC-MS, to determine the molecular fingerprints of waste degradation (Villas-Bôas et al., 2006). Our statistical analyses revealed subsets of active microorganisms highly associated to dynamics of substrate degradation over time, and enabled to posit novel hypotheses regarding the capacity of microorganisms to degrade specific metabolites.

## Methods

### Feedstock preparation

The inoculum used in the digestion experiments was sampled from a mesophilic full scale industrial anaerobic bioreactor treating primary sludge from a wastewater treatment plant (Valenton, France). The inoculum was incubated in anaerobic condition at 35°C without feeding during 10 days in order to degrade the organic matter in excess before carrying out the experiments.

Substrates used in the experiments were wastewater sludge collected from an industrial wastewater treatment plant (Valenton, France), fish waste obtained from a fish shop, and garden grass mowed from INRAE institute. Fish waste and garden grass were crushed and kept at 4°C before carrying out the experiments.

### Bioreactors experimental set-up

Binary mixtures of sludge with linearly decreasing percentages of either fish or grass were prepared (0-100, 25-75, 50-50, 75-25, 100-0 respectively, see Figure 1). Experiments were carried out in 1L glass bottles (700 mL working volume) at 35°C in the dark without agitation. The same quantity of carbon was added in all the bioreactors, and the ratio of substrate/inoculum used to feed and inoculate all the bioreactors was fixed at 12 gCOD/1.2 gCOD (Table S1). All bioreactors were complemented with a biochemical potential buffer (International Standard ISO 11734 (1995)) to reach a final working volume of 700 mL. All incubations were performed in triplicate. The bioreactors were then sealed with a screw cap and a rubber septum and the headspaces were flushed with N_2_ (purity >99.99 %, Linde gas SA). In total 27 anaerobic bioreactors were set-up.

**Figure 1.**
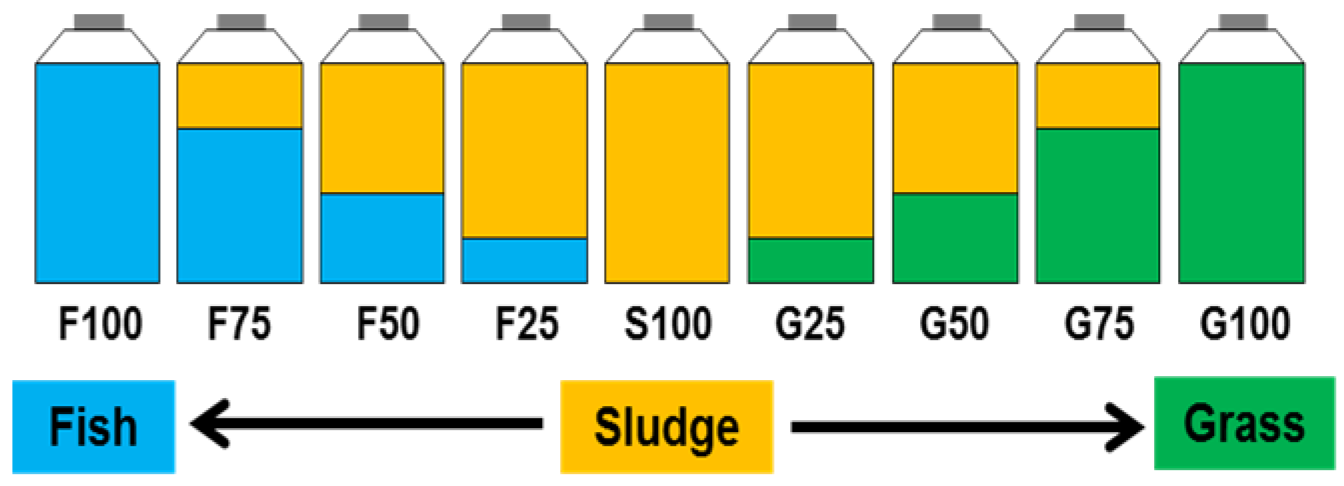
Scheme of the batch experimental design. S100 stands for bioreactors fed with wastewater sludge alone, F25, F50, F75, F100 stands for bioreactors fed with 25, 50, 75 or 100 % of fish (F) respectively, in co-digestion with sludge. G25, G50, G75, G100 stands for bioreactors fed with 25, 50, 75 or 100 % of Grass (G) respectively, in co-digestion with sludge.

In every reactor, 6 mL of liquid phase were sampled through the septum using a syringe at a weekly frequency (days 0, 14, 21, 28). The collected samples were centrifuged at 10 000g for 10 minutes to separate the supernatants from the pellets. Supernatants and pellets were snap frozen using liquid nitrogen. Supernatants were kept at −20°C for metabolomics analysis and pellets kept at −80°C for microbial analysis.

### RNA extraction and 16S rRNA sequencing

Based on the biogas production, a total of 22 samples were selected (corresponding to the highest biogas production as indicated in Figure 2). Since fish-fed bioreactors showed a higher delay than grass-fed bioreactors before biogas production started, the sampling time-points used were different for the two substrates (days 14 and 21 for grass-fed bioreactors, and days 21 and 28 for fish-fed bioreactors, Figure 2).

**Figure 2.**
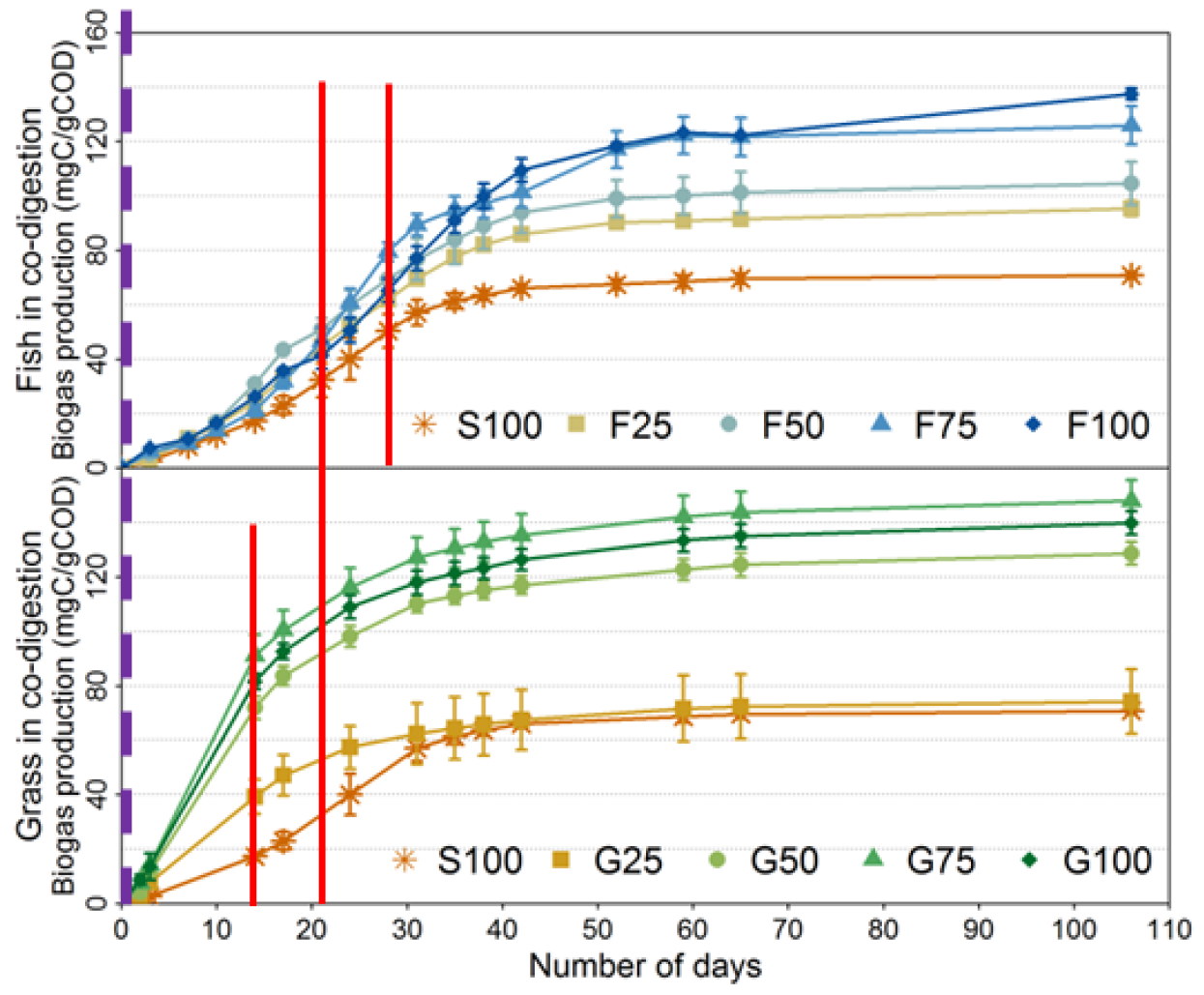
Cumulated biogas production (mgC/gCOD) over time (Days) for the different bioreactors. Mean values of the triplicate bioreactors are indicated with error bars representing standard deviations within triplicates. Co-digestion percentage is described in Figure 1, with either Fish (top) or Grass (bottom) co-digestion. Red solid lines indicate 16S rRNA sequencing and metabolomics sampling dates. Purple dashed line corresponds to the start of the incubation where an additional metabolomics analysis was carried out.

The commercial kit FastRNA Pro™ Soil-Direct (MP Biomedicals) was used to extract the total RNA following the manufacturer’s specifications. TURBO™ DNAse (Ambion) kit following the manufacturer’s instructions allowed to remove DNA co-extracted. The RNA was denaturated by 2 min at 85°C in a dry bath and was then stored on ice. RNAClean XP magnetic beads purification system (Beckman Coulter) was used to RNA purification by adding 1.8 volumes of beads by volume of RNA. After mixing by pipetting and 5 min of incubation, beads were captured using a magnetic rack on one side of the tube and then washed by adding 500 μL of 70% cold ethanol (diluted in DEPC-water). Tubes were incubated during 30 seconds at room temperature and ethanol was then removed. This washing step was repeated 3 times. Once ethanol finally evaporated, beads were resuspended with DEPC-water to eluted RNA from the beads. Finally, beads were removed using the magnetic rack and RNA was recovered in the supernatant. The integrity and quantity of the RNA was evaluated using High Sensitivity RNA ScreenTape and 2200 TapeStation (Agilent Technologies) following the manufacturer’s protocol.

A reverse transcription polymerase chain reaction (RT-PCR) was carried out on the RNA using the mix iScript Reverse Transcription Supermix (Biorad) and the following thermocycler program: 5 min at 25°C, 30 min at 42°C and 5 min at 85°C. The cDNA was quantified using Qubit 2.0 fluorometer (ssDNA assay kit, Invitrogen, Life Technologies).

Archaeal and bacterial hyper variable region V4-V5 of the 16S rRNA gene were amplified as cDNA, and these amplicons were then sequenced according to the protocol described by Madigou *et al.* (Madigou et al., 2019). The raw sequences were deposited in the NCBI databases under the project accession number PRJNA562430.

FROGS (Find Rapidly OTU with Galaxy Solution), a galaxy/CLI workflow (Escudié et al., 2018), was used to generate an OTU count matrix. R CRAN software (version 3.5.1) was used for alpha diversity analysis with Shannon method using phyloseq R package (version 1.20.0). Proportions of archaeal and bacterial OTUs abundances were scaled per sample and OTUs that exceeded 1% in terms of relative abundance in at least one sample, were selected for the remainder of the analysis and square-root transformed.

### Metabolomics analysis

Metabolomics analysis was performed on all collected supernatants. Samples were analysed using reverse phase liquid chromatography coupled to high resolution mass spectrometry (HPLC-ESI-HRMS) using a LTQ-Orbitrap XL instrument (Thermo Scientific). Samples were diluted at 1/10 in MilliQ water and 10 μL of the solution was injected into the analytical system. Chromatographic separation was performed on Accela 1250 pump at 400 μL/min with a linear gradient from 10 to 80 % of mobile phase A (acetonitrile + 0.05 % formic acid) and 90 to 20% of mobile phase B (water + 0.05% formic acid) into a Syncronis™ C18 column (50×2.1 mm, 1.7μm, Thermo Scientific) during 23 minutes, followed by a stabilisation phase of 5 minutes to return at the initial condition. After chromatographic separation, the sample was ionized by electrospray ionization (ESI) on positive mode. The detection was performed in full scan over an *m/z* range from 50 to 500 at a resolution of 100 000. A sample consisting of the supernatant from the digestion of anaerobic sludge was used as a quality control and injected every 10 experiment samples. Blank samples were injected every 10 samples, and an equimolar mix of the samples was injected every 5 samples.

The raw data obtained from the LC-HRMS analyses were transformed into mzXML files using MSConvert (ProteoWizard 3.0). The XCMS R package (version 1.52.0) was used to process the data (Smith et al., 2006). The method *centWave* was used to determine chromatographic peaks (region of interest - ROIs) with a *m/z* error of 10 ppm and a peakwidth between 20 and 50 seconds. The ROIs found in different samples were grouped using the group method with a bandwidth of 30. Retention times from the same ROI groups were unified across samples using the *orbiwarp* method. A second grouping was carried out using a bandwidth of 25. Finally, missing ROIs in the samples were filled using the *fillPeaks* method. Initial metabolite identification was performed based on the comparison of the accurate molecular mass measured by LC-HRMS with the corresponding values found in the online databases HMDB, LipidMaps, and PubChem (Fahy et al., 2007; Lee et al., 2018; Wishart et al., 2018). In addition, for selected compounds, a confirmation of the metabolite identification was performed by MS/MS fragmentation, and followed by the comparison of the acquired MS/MS spectra with the theoretical spectra from the online databases HMDB and MassBank (Horai et al., 2010).

### Statistical analysis and data integration

Multivariate data analysis was carried out using the mixOmics R package’ using a suite of component-based methods to identify groups of microorganisms and metabolites degradation with coordinated patterns.

First, each dataset was analysed independently using an exploratory (unsupervised) approach, or a supervised approach to identify key OTUs and metabolites prior to data integration as follows.

#### Unsupervised analysis on the 16S data

Sparse Principal Component Analysis (sPCA, (Shen and Huang, 2008)) was applied on all the samples of the microbial dataset (16S rRNA, 22 samples) to evidence main sources of variation and identify OTUs that most contribute to each principal component. Briefly, sPCA is based on PCA that includes a LASSO penalty for variable selection. In our context, the main source of variation correspond to specific feeding conditions (see Figure 3), and thus variable selection from sPCA enables us to identify microorganisms characteristic of co-substrate.

**Figure 3.**
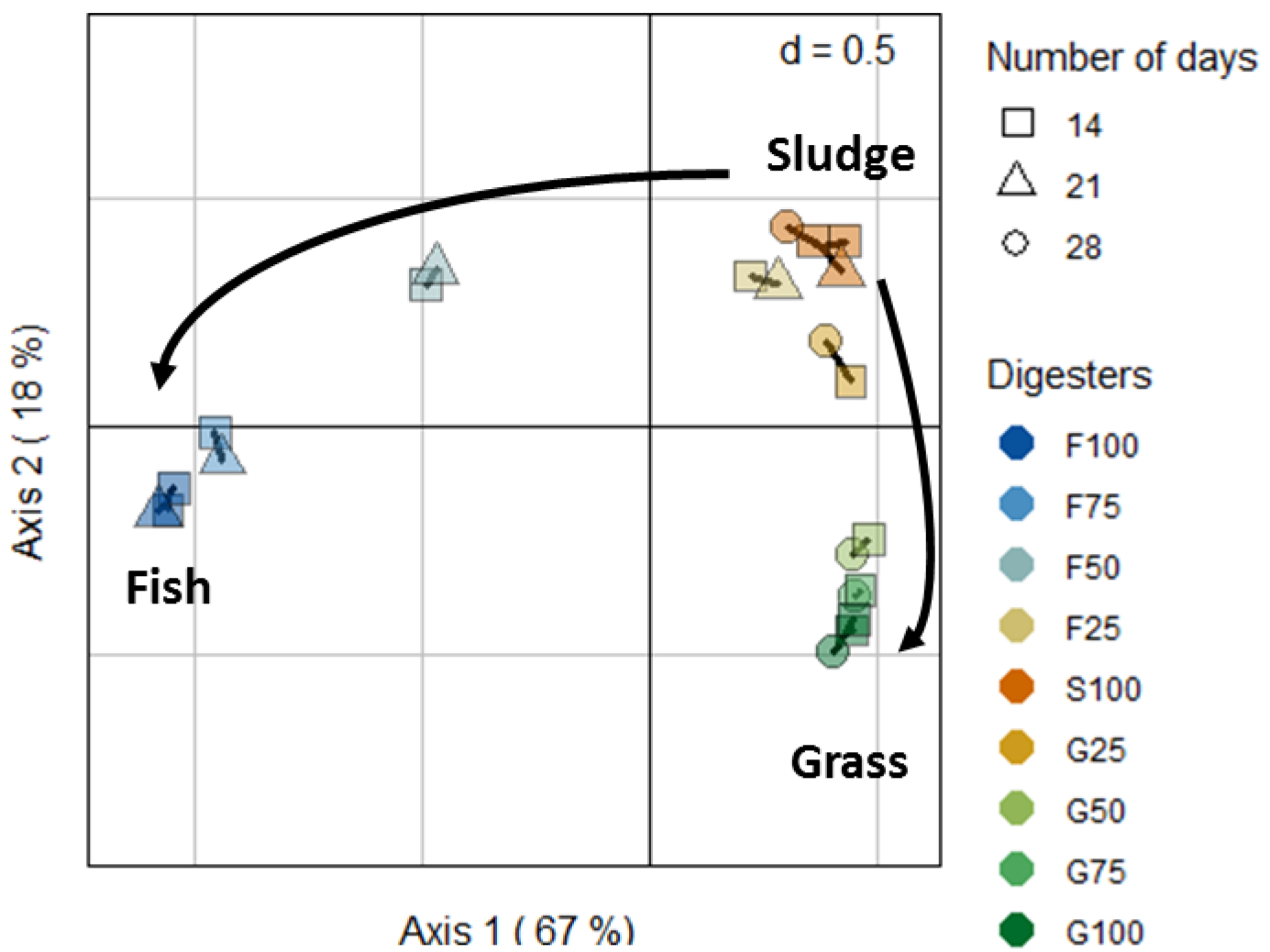
Microbial dynamics over time with respect to the different feeding compositions. Sample plot from the sPCA from the 16S rRNA dataset. Bioreactors are represented by colours and number of days by symbols.

#### Supervised analysis on the metabolomic day 0 data

Sparse Partial Least Squares Discriminant Analysis (sPLS-DA, (Lê Cao et al., 2011)), a supervised classification method, was applied on the mono-digestion samples (S100, F100, G100, 9 samples in total) at day 0 in order to determine the molecules specific to each substrate before any degradation. PLS-DA classifies samples into groups (sludge, fish, grass) and the sparse variant identifies metabolites discriminating each co-substrate at the start of the experiment. This method has already been used in metabolomics data analysis and successful in removing noise by selecting the most informative variables (e.g. Jiang et al., 2014). Classification performance was assessed based on the mean classification error rate using leave-one-out cross-validation, and Mahalanobis distance for prediction of the class of the samples (Rohart et al., 2017).

#### Clustering

Once selected, the abundances of the OTUs and the metabolites selected with sPCA and sPLS-DA respectively were clustered using hierarchical clustering (using Ward’s method and Manhattan and Euclidean distances respectively for each data type), then represented in heatmaps (heatmap.2 function from gplots R package, version 3.0.1).

#### Data integration of the selected OTUs and metabolites

The aim of this analysis is to identify patterns of different variables (microbes and metabolites) characteristics of the different conditions. Thus, this analysis included all 22 samples representative of the different feeding conditions. We calculated the metabolite degradation rate data as the ratio between the metabolites abundances at days 14, 21 or 28, relative to day 0. Both datasets were then integrated using Partial Least Square (PLS) regression method with a canonical mode (Lê Cao et al., 2009) to identify groups of OTUs and degraded metabolites that are highly correlated. Briefly, PLS weights variables from both datasets optimally using loading vectors, so that their linear combination (called PLS components) is maximally correlated and thus extract similar patterns across datasets (see supplemental figure S2 A and B). Loading vectors correspond to the weight assigned to each variable to define the PLS components. We subsequently applied hierarchical clustering on the loadings vectors to identify clusters of metabolites and OTUs variables with similar patterns of degradation rate or activity across the different types of substrate mixtures. As the correlation structure between high-throughput datasets can be highly spurious, we then further assessed the strength of the association between the variables assigned to each cluster using the proportionality distance (R package *propr* (Quinn et al., 2017)). We compared the distance between the profiles of each variable within one cluster and to the distance between the profiles outside this cluster. A small distance between two variables indicates a strong association.

## Results and discussion

### Influence of the feeding composition on the biogas production

We examined the cumulated biogas production in the different mixtures. Three main results can be observed from Figure 2. Firstly, the sludge mono-digestion produced a lower biogas quantity compared to the fish and grass mono-digestion. Secondly, the grass mono-digestion provided the best biogas production performances, as the biogas production rate and the final biogas production were higher than in fish mono-digestion. Thirdly, compared to sludge mono-digestion, the biogas production in bioreactors with mixtures of waste presented two patterns of improvement according to the type of co-substrate (fish or grass). In co-digestion with fish, the biogas production from the sludge improved linearly with the increase of fish quantity in the feedstock composition. In co-digestion with grass, the final biogas production was improved only when at least 50% of grass was present in the feedstock composition. A comparison of the major chemical parameters of anaerobic digestion has been already described in a precedent study (Cardona et al., 2019). In the present study, we focus on the influence of the feedstock composition on microbial activity and its relation with degradation of specific metabolites.

### Influence of the feeding composition on the microbial community

The influence of the feeding composition on the microbial diversity was evaluated using the Shannon index calculated on archaeal and bacterial communities (supplemental Figure S1). The bacterial diversity was inversely proportional to the amount of fish present in the bioreactors, while the level of the bacterial diversity remained stable when grass was present in co-digestion with sludge. The archaeal diversity decreased sharply with the presence of fish in the feedstock, or with high amount of grass. The low microbial diversity induced by the presence of fish can be explained by the composition of a simpler substrate (in terms of molecular variety) compared to the other substrates, or by a lower functional redundancy of the microorganisms that can degrade fish substrate compared to those that can degrade grass or sewage sludge.

Sparse PCA highlighted the main sources of variability in the 16S rRNA data set, while identifying key OTUs driving this variation. We selected 43 OTUs, which in combination explained 67 (first component) to 85% (two components) of the total microbial community variance in the data. The sample plot in Figure 3 showed the strong influence of the feeding composition on the microbial community. The samples were regrouped according to the major co-substrate, and regardless of the sample collection time. This result suggests that the variance due to time within conditions is low, and active microbial community over time is relatively stable.

The relative abundance of the selected microorganisms was compared between the different mixtures (Figure 4). We observed differences in the active archaeal community depending on the feeding type. Fish mono-digestion (F100) was mostly driven by the archaea *Methanosarcina* OTU_6 and *Methanoculleus* OTU_18; grass mono-digestion (G100) by *Methanospirullum* OTU_24*, Methanosarcina* OTU_1, *Methanofollis* OTU_64 and sludge mono-digestion (S100) by *Methanosarcina* OTU_1, *Methanobacterium* OTU_63 and 204 and two OTUs of *Methanoculleus* (18 and 50). With the exception of *Methanosarcina* which can use different methanogenesis pathways, all the identified archaea only use the hydrogenotrophic pathway to produce methane. The archaea composition is likely to depend on the environmental conditions in the bioreactors induced by the different feeding conditions and from bacterial dynamics. The fish substrate induced the activity of a specific OTU of *Methanosarcina* that was not found in the other substrates. As *Methanosarcina* presents a versatile methanogenesis metabolism, the modification of the OTU between the substrate could reflect the use of different methanogenesis metabolism.

**Figure 4.**
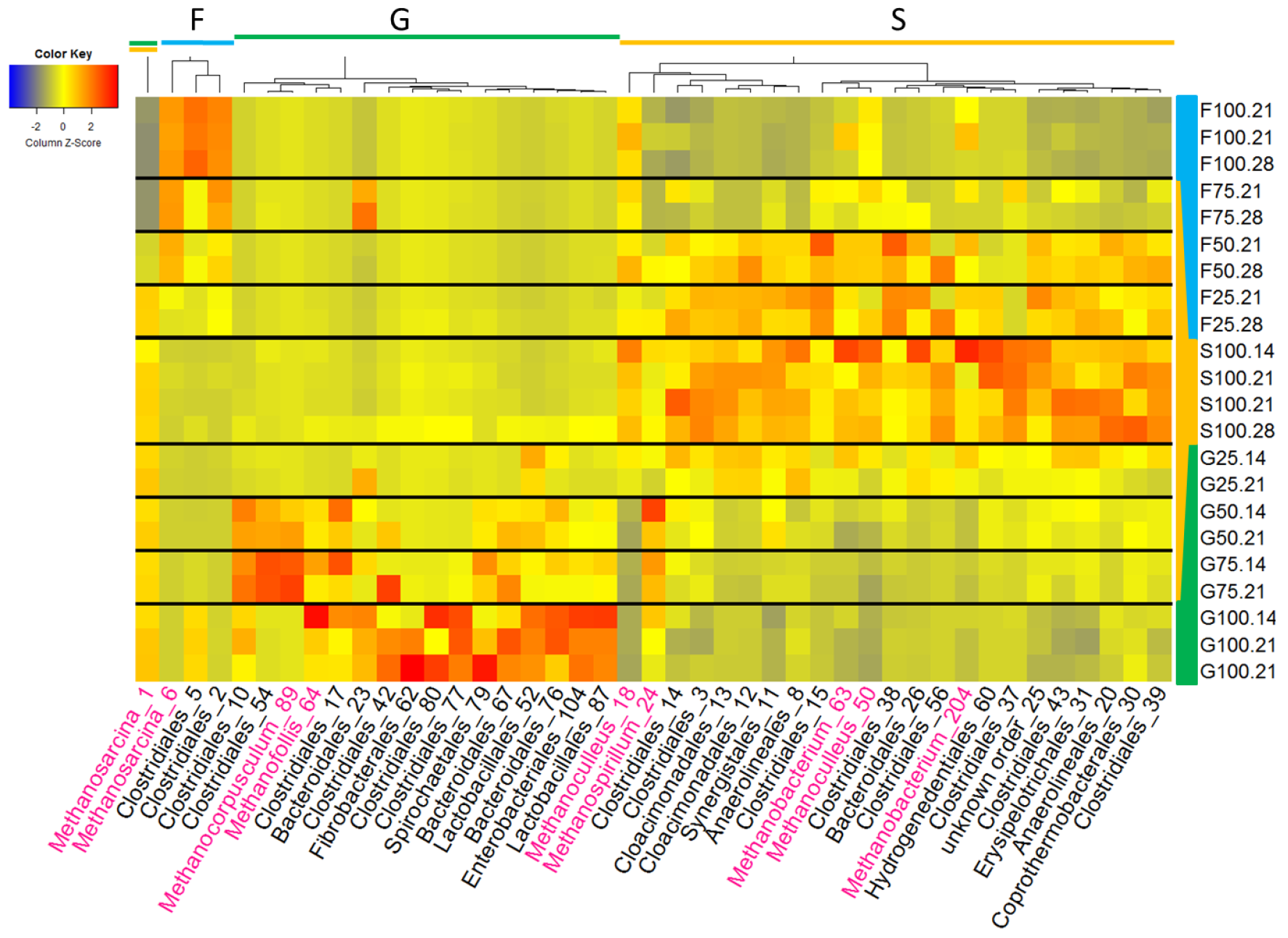
Heatmap of the microorganisms whose abundance contributes most to variance due to feeding composition selected with sPCA. The type of substrate used, day, duplicates were carried out on the bioreactors containing only fish, grass or sludge at day 21. Taxonomy completed by the OTU number is indicated at the genus level for archaea (pink) and at the order level for bacteria (black). Abundance is represented in the heatmap ranging from blue (low) to high (red).

The diversity in the bacteria community also differed between feeding types. As observed in the alpha diversity analysis, the fish substrate induced a lower microbial diversity than grass or sludge. In fish mono-digestion, the order *Clostridiales*, grouping cellulolytic degraders and fermentative members, represented more than 90% of the bacterial community while the abundance reached 25% and 50% for respectively sludge and grass mono-digestion. In grass mono-digestion, the abundance of *Spirochaetales*, *Fibrobacterales, Lactobacillales* and *Enterobacteriales* was higher than in fish and sludge mono-digestion. They were probably favoured by their ability to degrade cellulolytic substrates such as grass. Bacterial community of sludge mono-digestion was mostly composed by *Cloacimonadales*, *Synergistales*, *Anaerolineales*, *Hydrogenedentiales*, *Erysipelotrichales* and *Coprothermobateriales*. Members from the orders *Synergistales, Cloacimonadales, Coprothermobacteriaceae* and *Anaerolineales* are known or suspected to be able to form syntrophic interaction with hydrogenotrophic methanogens (Calusinska et al., 2018; Gagliano et al., 2015; Ito et al., 2011; Sekiguchi et al., 2001). *Hydrogenedentiales* are H_2_-utilising bacteria and associations with the H_2_-producers *Coprothermobacteriaceae* were evidenced (Nobu et al., 2015). In sludge, the abundance of potential syntrophic partners in presence with hydrogenotrophic methanogens suggests that methane production was mainly produced from the hydrogenotrophic methanogenesis pathway, in line with the archaeal community described in this incubation.

In light of the current literature, the type of substrate used to feed the bioreactors has been suggested to contribute to the development of an adapted microbial community of degraders (De Francisci et al., 2015; Lee et al., 2018). Specifically, in our study, fish substrate was found to induce not only a low microbial diversity, but also the growth of a specific community compared to what was observed in grass and in sludge substrates. Indeed, only 5% of the selected OTUs were common between fish- and sludge-bioreactors, 9% between both fish- and grass-bioreactors, while 17% of the selected OTUs were common between both grass- and sludge-bioreactors.

Within the mixtures, the abundance of the selected microorganisms changed with feeding composition. However, microbial activity did not necessarily increase with the amount of mixture proportions. For example, the relative abundances of the OTUs identified as active in the fish:sludge mixture at 75:25 (F75) were similar to the relative abundances of the same active OTUs in the fish mono-digestion (F100). As the proportion of sludge increased, the active microorganisms more characteristic of sludge increased as well. However, in grass mixtures, the active microbial community from sludge remained dominant regardless the proportion of grass used (e.g down to 25% in the mixture with 75% of sludge, G25). However, some microorganisms were found in specific proportions of grass:sludge. For example, *Methanocorpusculum* and specific OTUs of *Clostridiales* were found in the mixes 50:50 and 75:25 of grass:sludge.

### Substrates degradation dynamics

We studied the temporal dynamics of degradation of the different substrate mixtures by analysing the metabolic fingerprint of the degradation in the bioreactors using an HPLC-ESI-HRMS instrument. After data examination with XCMS, a total of 267 regions of interest (ROI) were detected. ROI designate chromatographic features (m/z) in the spectrum. Sparse PLS-DA identified 70 ROIs out of the 267 ROIs specific to each substrate (sludge, grass, fish) initially present at day 0. The classification error rate from the sPLS-DA was 0, indicating that all samples could be perfectly assigned to their respective groups using leave-one-out cross validation based on these 70 ROIs. The ROIs were then clustered into four groups according to their intensities at day 0. The degradation pattern of the selected ROIs was then examined for every mixture across all time points. Four main degradation patterns were observed (Figure 5). Three of these groups include the ROIs whose relative intensities were high only in sludge-, grass- or fish-fed bioreactors (indicated as S, G, and F, respectively, in Figure 5) at day 0. The other group (G and F) include the ROIs whose relative intensities were high in both fish- and grass-but low in sludge-containing bioreactors at day 0.

**Figure 5.**
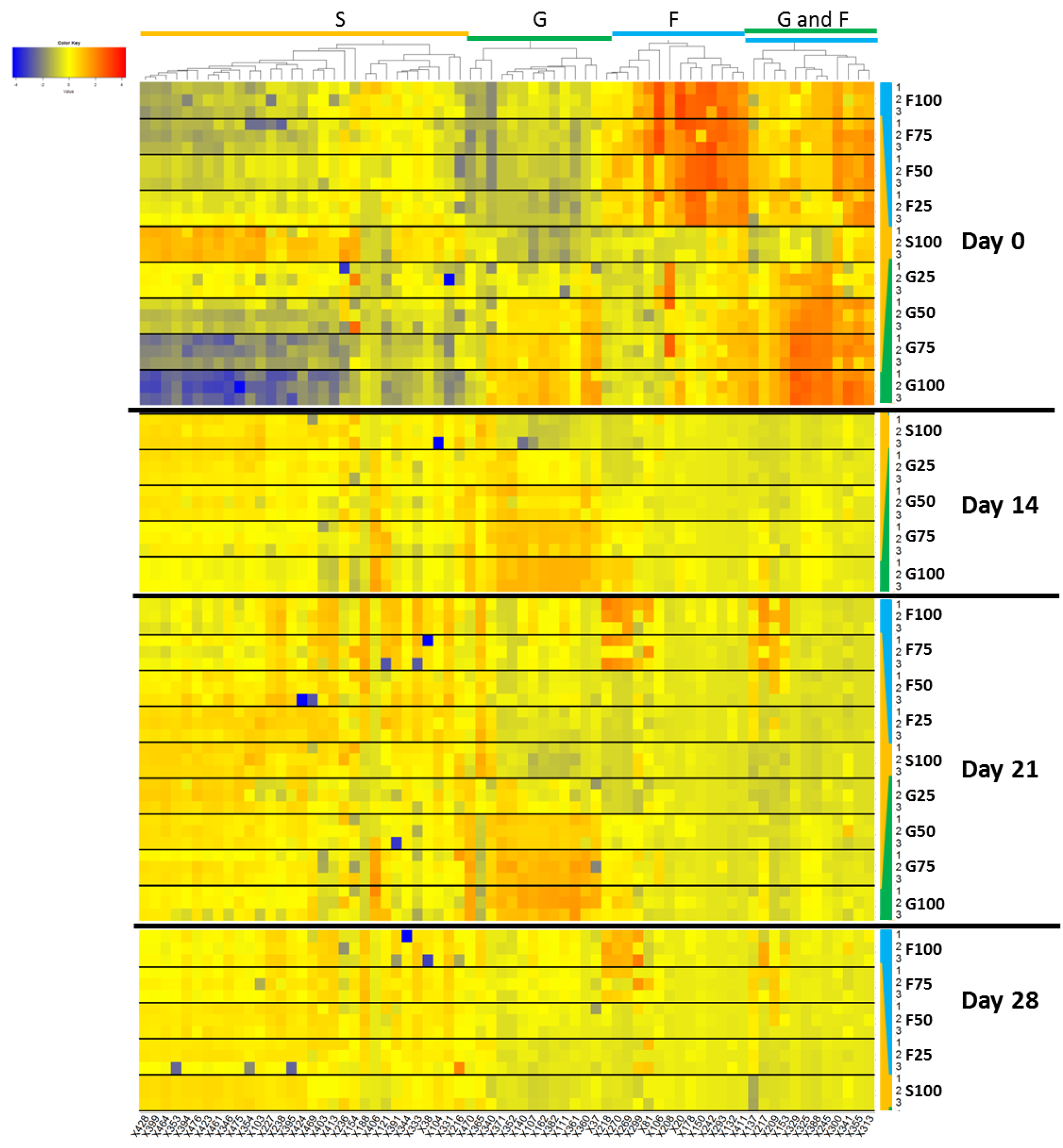
Metabolites dynamics within bioreactors and over time. The most discriminant ROIs for the different co-substrates were selected by sPLS-DA at day 0, then clustered with hierarchical clustering according to day 0. The profiles for the remaining days were then represented in the heatmap based on the intensities. The different panels represent days 21-28 for bioreactors containing fish, 14-21 for bioreactors containing grass and 14-21-28 for bioreactors containing sludge. For each waste mixture and date, triplicates bioreactors numbered 1, 2 and 3 are indicated. Intensity ranges from blue (low) to red (high).

Whilst the ROIs were selected to discriminate only mono-digestion at day 0, we observed that the intensity of the ROIs differed across all feeding types. This result was expected given the differences in the molecular composition of the substrates. In general, the intensity of the ROIs representative of sludge decreased when fish or grass was mixed with sludge. We also observed some temporal dynamics, as that the intensity of some ROIs decreased over time, while some other either increased or remained stable during the experiment. This can be explained as some metabolites may not be easily degraded, or may be a product of degradation of other metabolites.

The putative identification of the ROIs based on their *m/z* is given in the Table S2. Out of the 70 selected ROIs, 19 metabolites were identified, whilst the rest could not be assigned. The metabolites assignment step in MS metabolomics analysis is a traditional bottleneck (Longnecker et al., 2015). Our study was further hampered by the high structural diversity of the compounds found in bioreactors, the lack of databases specific for anaerobic digesters, and the absence of relevant literature of metabolomics analysis on anaerobic bioreactors. However, the metabolites identified in the different clusters were biologically consistent with the corresponding substrates. For example, metabolites identified in sludge-containing samples were diethylthiophosphate (X340) and 6-methylquinoline (X218 and X270). Diethylthiophosphate is a common degradation product of organophosphorus pesticides, while 6-methylquinoline is a flavouring ingredient found in tea. Metabolites characteristics of grass-fed bioreactors were plant constituents (betaine, X245), metabolites obtained from lignin degradation (trans-ferulic acid and p-coumaric acid, X388 and X365, respectively), and metabolites from sugar metabolism (galactitol, X329). In fish-fed bioreactors, most of the metabolites corresponded to organic compounds resulting from amino acids degradation (cadaverine and histamine, X132 and X150, respectively).

### Correlation between microbial activity and substrates degradation pattern

Statistical data integration analysis was carried out between the microbial and metabolites datasets to identify potential degraders of specific metabolites. As far as we know, this type of analysis has never been applied in the anaerobic digestion field. Correlation studies have been used to relate metabolite profiles to microbiome in gut studies (Han et al., 2019; McHardy et al., 2013). For example, Han et al (2019) estimated the correlation between cecal compounds and bacteria profiles acquired respectively by LC/MS and 16S RNA sequencing analyses using Spearman’s correlation. However, in datasets containing a large number of variables, such approach is highly likely to result in spurious correlations. One novelty in our study is to use component-based approaches, such as multivariate PLS method that are well suited for a high correlation structure within and between datasets, whilst being able to highlight key associations between variables from each dataset.

As described in the methods section, we considered the metabolites degradation rates relative to day 0 in our analysis. The sample (ordination) plots from PLS (Figure S2 A-C) show similar patterns for microbial and metabolites datasets, suggesting that an underlying correlation structure between the two datasets exists and can be extract with such method. The correlation circle plot (Figure S2 D) projects the variables into the space spanned by the PLS components to visualise simultaneously the active microorganisms and the metabolites responsible for this ordination of the samples (González et al., 2012). In particular, active microorganisms and metabolites with a high degradation rate projected close to each other on this graph are correlated, as they contribute equally to define the PLS components. A hierarchical clustering was performed on the PLS loadings of the microbial and metabolites variables. It helped to further identify groups of microorganisms potentially associated with the metabolites degradation. In total, five clusters were evidenced.

To illustrate the associations, the Figure 6 depicts the mean values of the microbial activity and metabolite degradation rate according to the feeding types at day 21. Microorganisms and metabolites showing the same evolution in the different feeding mixtures are clustered together. Similar trends were observed at the other dates (see supplemental figure S4). The quality of the clustering was further assessed by comparing the proportionality distances between profiles (Bodein et al., 2019; Quinn et al., 2017). The distances were lower between profiles grouped within the same cluster, compared to the distances with the profiles assigned to other clusters (supplemental Figure S3 and Table S3). This result confirmed a strong association between the profiles of the active microorganisms and metabolites across all feeding types, and their links where further investigated.

**Figure 6.**
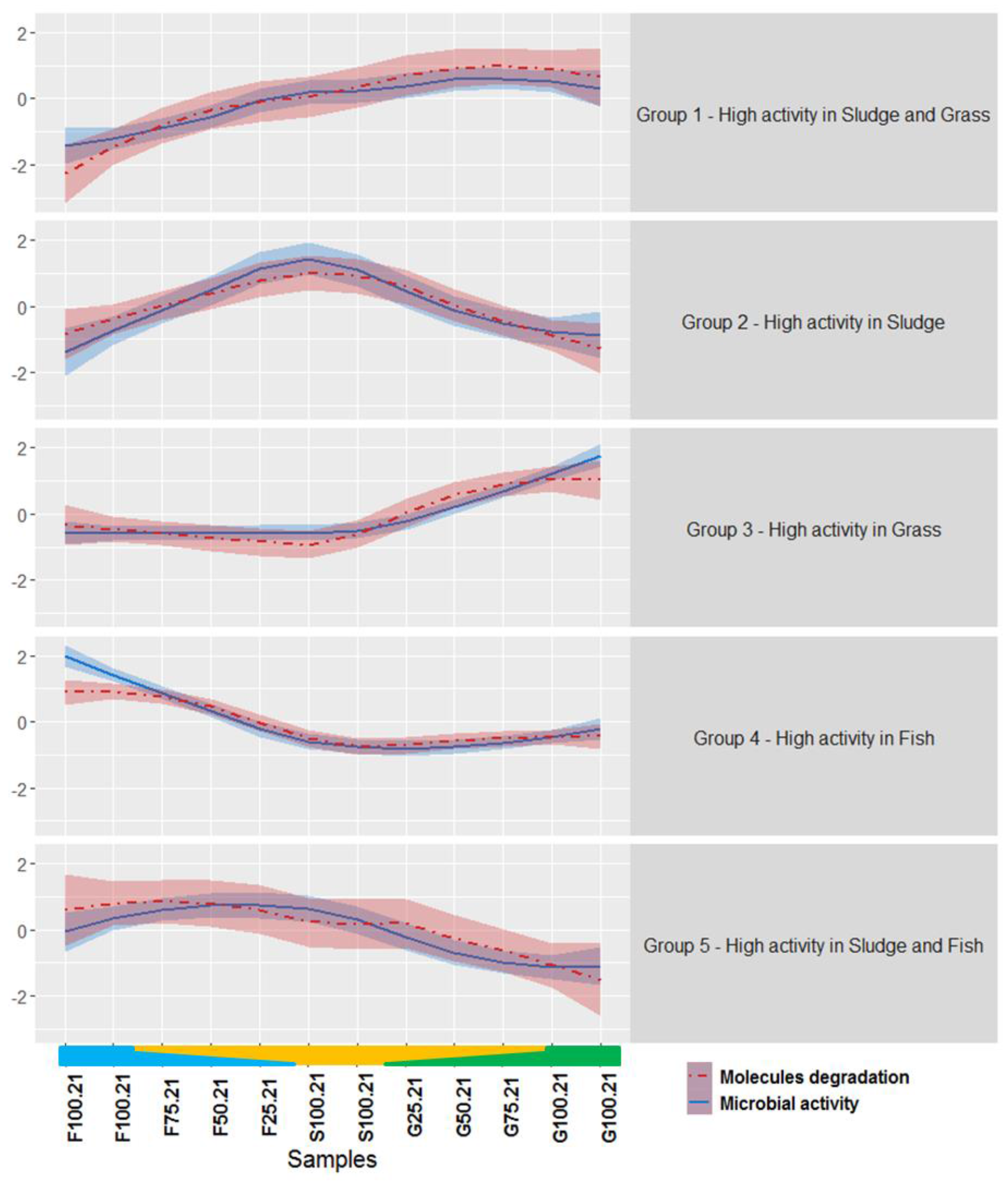
Correlated dynamics of the active microorganisms and metabolites degradation across all samples at day 21. Lines represent the mean values of the different microbial activity (solid blue line) or metabolites degradation rates (dashed red line) for each of the clusters identified using hierarchical clustering performed on the PLS loadings. Shadows represent standard deviation. S100 stands for wastewater sludge alone, F25, F50, F75, F100 stands for respectively 25, 50, 75 or 100% of fish (F) in co-digestion with sludge, G25, G50, G75, G100 stands for respectively 25, 50, 75 or 100% of Grass (G) in co-digestion with sludge.

The microorganisms and metabolites assigned to the five clusters are described in the supplemental Tables S3 and S4. Group 1 included microorganisms and metabolites with a high microbial activity and high metabolite degradation rate during the digestion of sludge and grass. Groups 2 to 4 included microorganisms and metabolites that were specific of either sludge, grass, or fish, respectively. Finally, group 5 included the microorganisms and metabolites that were highly active and highly degraded, respectively, in fish and sludge bioreactors.

Group 1 included two genera of archaea, *Methanosarcina* and *Methanospirullum*, and the metabolites diethylthiophosphate and N-(3-methylbutyl)acetamide (X153). As mentioned before, diethylthiophosphate is a pesticide degradation product found in urine (Nomura et al., 2014) while acetamides are produced in the fermentation of vegetal matter. One hypothesis explaining the association between these archaea and metabolites could be an indirect role of the archaea in the metabolites degradation through a syntrophic interaction with bacteria. This is the first time such association is reported, and further investigations are required.

Group 2 included 14 OTUs from the orders *Cloacimonadales*, *Clostridiales*, *Anaerolineales, Synergistales, Bacteroidales, Hydrogenedentiales* and *Coprothermobacterales*, and 8 metabolites including compounds from tryptophan degradation (L-tryptophanol and tryptamine, X217 and X269, respectively), 6-methylquinoline and thioxoacetic acid, among others. In agreement with these metabolites, *Anaerolineales* and *Synergistales* are known for their ability to degrade amino acids (Swiatczak et al., 2017). Thus, their correlation with L-tryptophanol and tryptamine is consistent with the literature. Surprisingly, no methanogen was identified in this group as we would have expected to find with these syntrophic bacteria. One reason could be that the methanogens were not specific partners of these bacteria. Indeed, most of the methanogens were found ubiquitously in bioreactors fed partly with either sludge or grass (groups 2 and 3) as shown in Figure 4.

In group 3, 12 OTUs mostly from the orders *Clostridiales*, *Lactobacillales*, *Bacteroidales* and *Spirochaetales* and the archaea *Methanofollis* were found associated with 7 metabolites from sugar degradation, (galactitol), lignin compounds (p-coumaric, trans-ferulic acid), plant constituent (betaine) or histidine degradation (Imidazolepropionic acid). From this group, it is worth noting the association between the microbial activity of the lactic acid bacteria *Lactobacillales* and the degradation of lignin compounds such as trans-ferulic acid (X388) and p-coumaric acid (X365). Indeed, such microorganisms are lignin degraders (Fessard and Remize, 2017; Filannino et al., 2014).

In group 4, *Methanosarcina* and 2 OTUs from the order *Clostridiales* and metabolites classified as amino acids degradation products were identified. Cadaverine (X132) and 5-aminopentanoic acid (X208) are obtained from L-lysine degradation, histamine (X150) from L-histidine degradation, and phenylpyruvic acid (X300) from L-phenylalanine degradation. The presumed role of these microorganisms in the degradation of cadaverine and L-histidine can be supported by previous studies. Roeder and Schink described a new strain close to *Clostridium aminobutyricum,* able to degrade cadaverine, in co-culture with the archaea *Methanospirullum* (Roeder and Schink, 2009). Some *Clostridium* were also identified to be involved in the histamine degradation (Pugin et al., 2017).

In Group 5, heptane-1,2,3-triol and hexadecandiperoxoic acid were found to be descriptive of this cluster. The origin of these metabolites is unclear and must be further investigated. *Methanoculleus* and *Methanobacterium* were correlated to the genus *Syntrophomonas*. Some species of this bacterium are known to growth in syntrophy with H_2_-consumer as methanogens (Mcinerney et al., 1981).

Thus, the refinement of our clusters has highlighted potentially relevant associations between molecules degradation and microbial activity across different feeding substrates. This is the first time that such extensive data integration analysis is performed in anaerobic digestion. Previous multi-omics studies only considered single omics analysis (Beale et al., 2016; Bize et al., 2015; Joyce et al., 2018).

Our analytical workflow suggests relationships between the degradation of metabolites by different microorganisms. These relationships were consistent with the literature and future work will be carried out to validate them. For instance, the degradation of the metabolites by the identified microorganisms could be confirmed with either microbial cultures using these specific metabolites as substrates, or by performing experiments with labelled metabolites, such as stable isotope probing (Chapleur et al., 2016).

## Conclusion

This study proposed a suite of statistical tools to unravel associations between molecules degradation and microbial activity, whilst carefully preventing spurious correlations between them. We applied component-based approaches for variable selection and data integration, then further evaluated the clusters of variables using proportionality distances. Our analyses show the existence of links between the anaerobic digester feeding composition and microbiota activity. The associations identified between the microorganisms’ activity and the degradation of metabolites were biologically relevant and consistent with previous literature. The statistical framework we developed in this study are promising for the the screening analysis of potential metabolites degraders in different complex ecosystems such as other bioprocesses, or human microbiota.

## Acknowledgements

We thank Nadine Derlet from the Irstea PROSE analytical division for her technical support. We acknowledge SUEZ Environment for providing us the access to the wastewater treatment plant of Valenton.

