## supplemental for "Integrative analyses to investigate the link between microbial activity and metabolites degradation during anaerobic digestion"

**Table S1. Mixture details for the different bioreactors.**

| Name | Substrate ratios (%) | Sludge (g) | Fish (g) | Grass (g) | Inoculum (g) |
| --- | --- | --- | --- | --- | --- |
| F100 | Fish | 0 | 39 | 0 | 93 |
| F75 | Fish:Sludge 75:25 | 29 | 29 | 0 | 93 |
| F50 | Fish:Sludge 50:50 | 58 | 20 | 0 | 93 |
| F25 | Fish:Sludge 25:75 | 87 | 10 | 0 | 93 |
| S100 | Sludge | 116 | 0 | 0 | 93 |
| G25 | Grass:Sludge 25:75 | 87 | 0 | 32 | 93 |
| G50 | Grass:Sludge 50:50 | 58 | 0 | 64 | 93 |
| G75 | Grass:Sludge 75:25 | 29 | 0 | 95 | 93 |
| G100 | Grass | 0 | 0 | 126 | 93 |

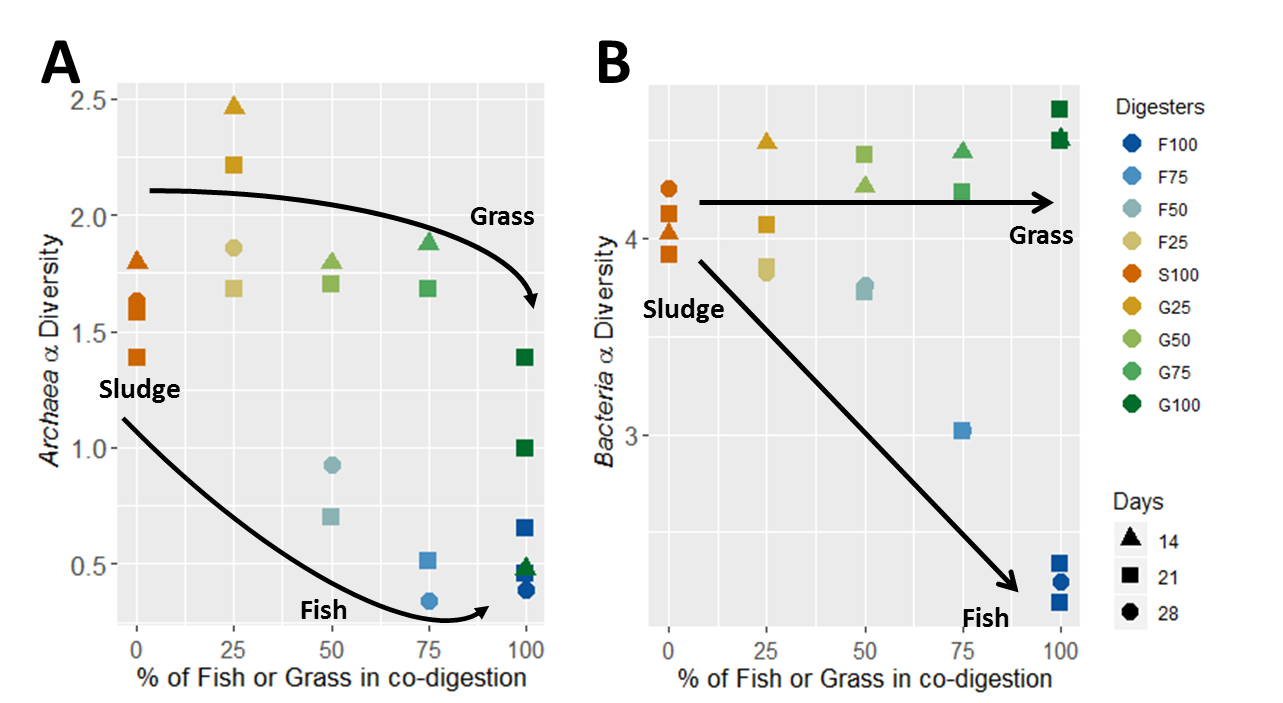

**Figure S1. Alpha diversity of the total active microbial community within the mixtures.** Shannon method was used to determine the archaea (A) and bacteria (B) alpha diversity for the different feeding composition at days 21-28 for bioreactors containing fish, 14-21 for bioreactors containing grass and 14-21-28 for bioreactors containing sludge only. Duplicate analyses were carried out in the bioreactors containing only fish, grass, or sludge at day 21.

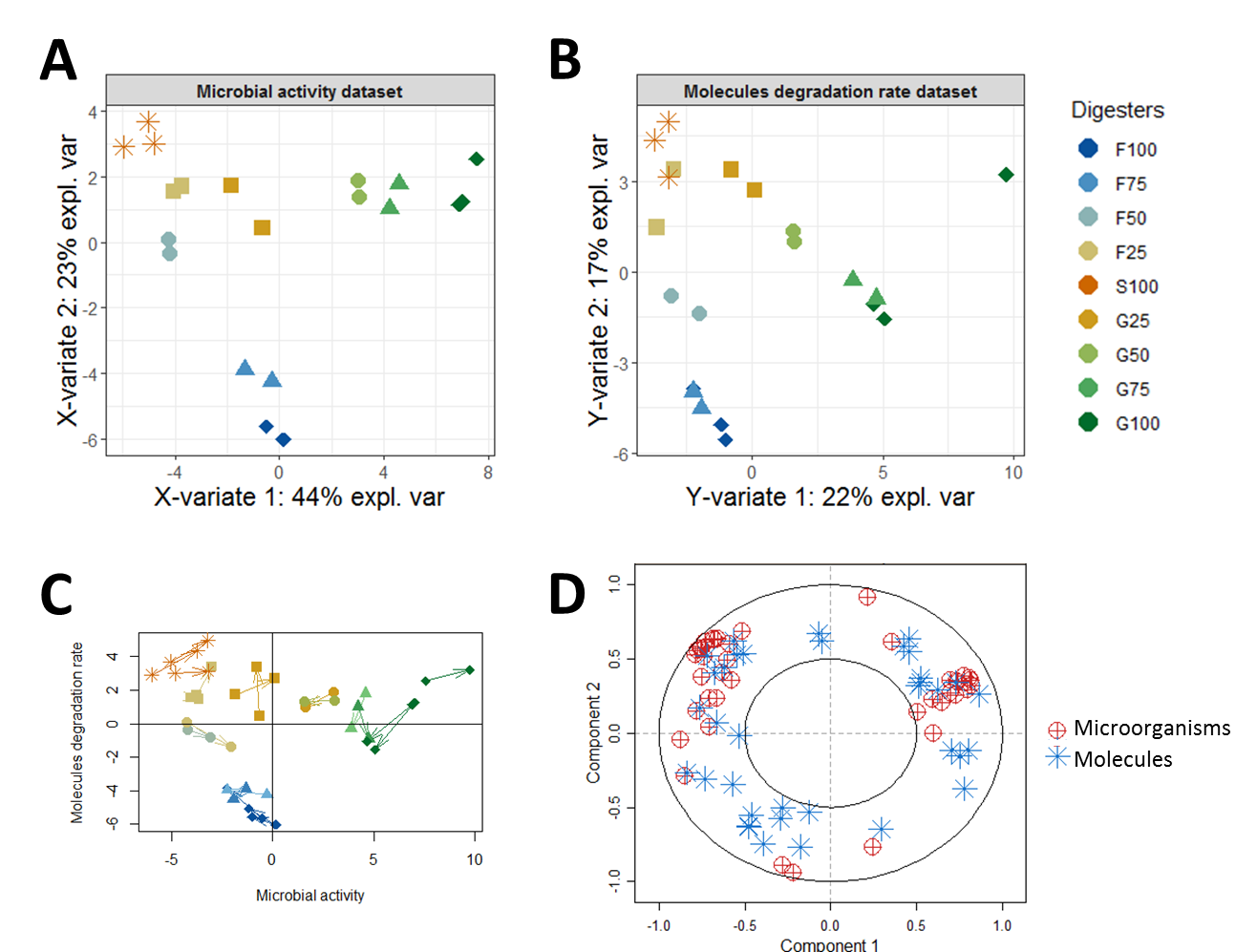

Figure S2. PLS data integration of the microbial activity and molecules degradation rate. Sample ordination plots according to the microbial activity (A) and molecules degradation dataset (B) show a similar influence of the feeding composition on the microorganisms and on the degradation of the molecules. In C, each arrow links match samples from the two superimposed ordination plots. Short arrows represent a strong similarity in the information contained in both datasets. The correlation circle plot (D) shows the microorganisms and the molecules with similar dynamics across the different samples.

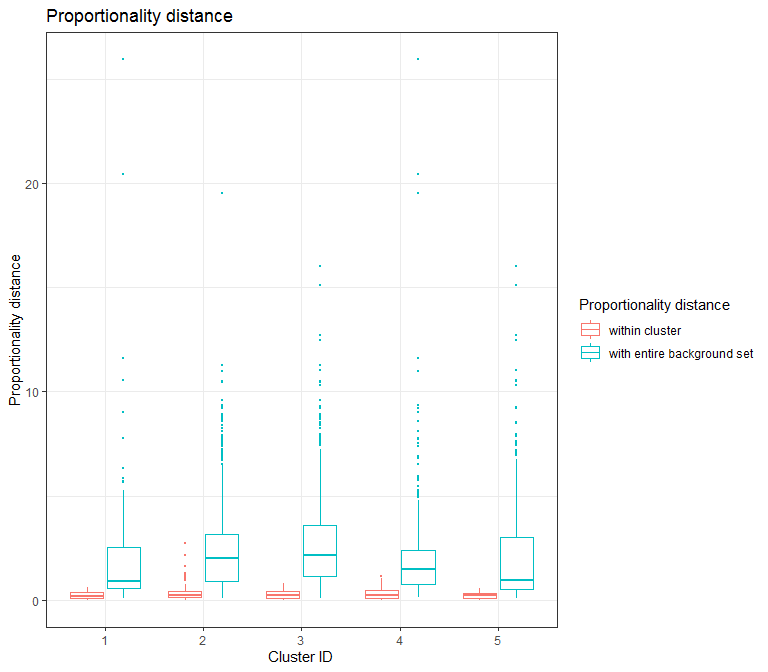

**Figure S3. Estimation of the proportionality distance per cluster identified with hierarchical clustering.** The distance was calculated between each profile within a cluster and with the residual profiles set (profile outside the cluster).

**Table S3: Proportionality distance per cluster identified with hierarchical clustering.** The median distance is reported for the profiles within and outside the cluster for each cluster. The Wilcoxon test pvalue was estimated to evaluate the difference between the median.

| **Cluster** | **Median inside** | **Median outside** | **Wilcoxon Test pvalue** |
| --- | --- | --- | --- |
| **1** | 0.21 | 0.91 | 1.34-8 |
| **2** | 0.26 | 2.03 | 9.20-166 |
| **3** | 0.23 | 2.17 | 3.93-152 |
| **4** | 0.23 | 1.46 | 3.68-58 |
| **5** | 0.25 | 0.98 | 2.33-26 |

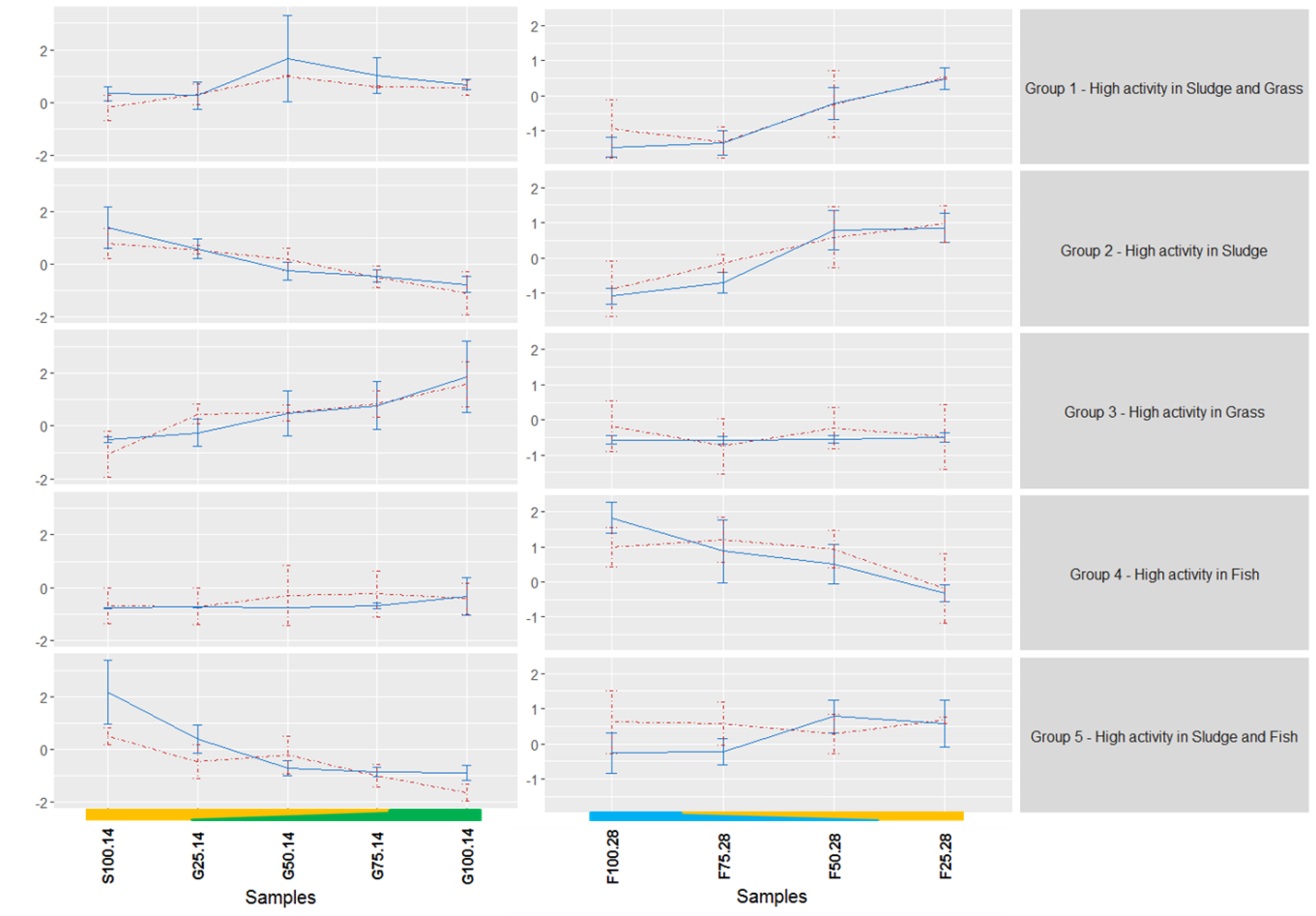

**Figure S4. Correlated dynamics of the active microorganisms and metabolites degradation across the samples at day 14 and day 28.** Lines represent the mean values of the different microbial activity (solid blue line) or metabolites degradation rates (dashed red line) for each of the clusters identified using hierarchical clustering performed on the PLS loadings. Shadows represent standard deviation. S100 stands for wastewater sludge alone, F25, F50, F75, F100 stands for respectively 25, 50, 75 or 100% of fish (F) in co-digestion with sludge, G25, G50, G75, G100 stands for respectively 25, 50, 75 or 100% of Grass (G) in co-digestion with sludge.

**Table S4: Putative identification of the molecules in each cluster**

| **Cluster** | **Substrate specificty** | **Feature** | **Measured mass (in Da)** | **Elemental composition** | **Calculated mass (in Da)** | **Error** | **Ion assignment** | **MS/MS Fragments** | **Molecule name** | **Databases Code** | | |
| --- | --- | --- | --- | --- | --- | --- | --- | --- | --- | --- | --- | --- |
|  |  |  |  |  |  | **(in ppm)** |  |  |  | **HMDB** | **PubChem** | **LipidMaps** |
| **1** | **Sludge or Grass** | X340 | 214.98765 | C_4_H_10_O_3_PSNa_2_ | 214.98782 | -0.7684 | [M+2Na-H]^~~+~~^ | ^1^n/a | Diethylthiophosphate | HMDB0001460 | 655 |  |
|  |  | X153 | 116.1069 | C_6_H_14_NO | 116.10699 | -0.7264 | [M+NH_4_]^+^ | ^1^n/a | N-(3-Methylbutyl)acetamide | HMDB0031651 | 98643 |  |
| **2** | **Sludge** | X346 | 218.91951 | C_4_H_4_O_4_S_2_K | 218.91881 | 3.2144 | [2M+K]^+^ | ^1^n/a | Thioxoacetic acid |  | 18612521 |  |
|  |  | X218 | 144.08073 | C_10_H_10_N | 144.08132 | -4.1567 | [M+H]^+^ | ^1^n/a | 6-Methylquinoline | HMDB0033115 | 7059 |  |
|  |  | X270 | 161.1072 | C_10_H_13_N_2_ | 161.10787 | -4.1705 | [M+NH_4_]^+^ | ^1^n/a | 6-Methylquinoline | HMDB0033115 | 7059 |  |
|  |  | X269 | 161.10723 | C_10_H_13_N_2_ | 161.10787 | -3.983 | [M+H]^+^ | 144.0804 | Tryptamine | HMDB0000303 | 1150 |  |
|  |  | X217 | 144.08073 | C_10_H_10_N | 144.08132 | -4.1567 | [M+H-H_2_O]^+^ | ^1^n/a | L-Tryptophanol | HMDB0003447 | 6951149 |  |
|  |  | X469 | 436.88109 |  |  |  |  | ^1^n/a |  |  |  |  |
|  |  | X464 | 430.91425 |  |  |  |  | ^1^n/a |  |  |  |  |
|  |  | X361 | 234.89398 |  |  |  |  | ^1^n/a |  |  |  |  |
| **3** | **Grass** | X365 | 240.96739 | C_9_H_7_O_3_K_2_ | 240.96694 | 1.8974 | [M+2K-H]^~~+~~^ | ^1^n/a | p-Coumaric acid | HMDB0002035 | 637542 |  |
|  |  | X245 | 156.04210 | C_5_H_11_NO_2_K | 156.04269 | -3.7528 | [M+K]^+^ | 112.0520 | Betaine | HMDB0000043 | 247 |  |
|  |  | X388 | 270.97801 | C_10_H_9_O_4_K_2_ | 270.97805 | -0.138 | [M+2K-H]^~~+~~^ | ^1^n/a | Trans-ferulic acid | HMDB0000954 | 445858 |  |
|  |  | X209 | 141.06586 | C_6_H_9_N_2_O_2_ | 141.06585 | 0.0226 | [M+H]^+^ | 123.0549 | Imidazolepropionic acid | HMDB0002271 | 70630 |  |
|  |  | X329 | 205.06843 | C_6_H_14_O_6_Na | 205.06881 | -1.8535 | [M+Na]^+^ | ^1^n/a | Galactitol | HMDB0000107 | 11850 |  |
|  |  | X137 | 105.11022 |  |  |  |  | ^1^n/a |  |  |  |  |
|  |  | X188 | 130.12247 |  |  |  |  | ^1^n/a |  |  |  |  |
| **4** | **Fish** | X208 | 140.06812 | C_5_H_11_NO_2_Na | 140.06875 | -4.5164 | [M+Na]^+^ | ^1^n/a | 5-Aminopentanoic acid | HMDB0003355 | 138 |  |
|  |  | X178 | 128.07052 | C_6_H_10_NO_2_ | 128.0706 | -0.6371 | [M+NH_4_]^+^ | ^1^n/a | 2.4-Hexadienedial |  | 5283312 | LMFA06000011 |
|  |  | X150 | 112.08674 | C_5_H_10_N_3_ | 112.08692 | -1.647 | [M+H]^+^ | 95.0601 | Histamine | HMDB0000870 | 774 |  |
|  |  | X132 | 103.12285 | C_5_H_15_N_2_ | 103.12297 | -1.2069 | [M+H]^+^ | 86.0961 | Cadaverine | HMDB0002322 | 273 |  |
|  |  | X300 | 182.08113 | C_9_H_12_NO_3_ | 182.08117 | -0.1928 | [M+NH_4_]^+^ | ^1^n/a | Phenylpyruvic acid | HMDB00205 | 997 |  |
|  |  | X81 | 89.11519 |  |  |  |  | ^1^n/a |  |  |  |  |
|  |  | X20 | 69.06968 |  |  |  |  | ^1^n/a |  |  |  |  |
|  |  | X371 | 245.13844 |  |  |  |  | ^1^n/a |  |  |  |  |
|  |  | X352 | 227.12705 |  |  |  |  | ^1^n/a |  |  |  |  |
| **5** | **Sludge or Fish** | X236 | 149.11705 | C_7_H_17_O_3_ | 149.11722 | -1.1307 | [M+H]^+^ | ^1^n/a | Heptane-1,2,3-triol |  | 124604 | LMFA05000665 |
|  |  | X406 | 301.20101 | C_16_H_29_O_5_ | 301.2015 | -1.6245 | [M+H-H_2_O]^+^ | ^1^n/a | Hexadecanediperoxoic acid |  | 22027452 |  |
|  |  | X424 | 354.89501 |  |  |  |  | ^1^n/a |  |  |  |  |

^1^n/a: MS/MS spectrum not acquired for the given ion.

**Table S5: Active microorganisms identified in each cluster**

| **Cluster** | **Feeding type** | **Microorganisms identification** | **Order and genus taxonomy information** |
| --- | --- | --- | --- |
| 1 | Sludge and Grass | Methanosarcina_1 | Methanosarcinaceae - Methanosarcina |
|  |  | Methanospirillum_24 | Methanospirillaceae - Methanospirillum |
| 2 | Sludge | Cloacimonadales_12 | Cloacimonadaceae - W5 |
|  |  | Cloacimonadales_13 | Cloacimonadaceae - W5 |
|  |  | Clostridiales_3 | Clostridiaceae 1 - Clostridium sensu stricto 1 |
|  |  | Anaerolineales_8 | Anaerolineaceae - Flexilinea |
|  |  | Synergistales_11 | Synergistaceae - unknown genus |
|  |  | Anaerolineales_20 | Anaerolineaceae - unknown genus |
|  |  | Clostridiales_14 | Peptostreptococcaceae - Multi-affiliation |
|  |  | unknown order_25 | BRC1 - unknown genus |
|  |  | Clostridiales_39 | Clostridiaceae 1 - unknown genus |
|  |  | Bacteroidales_60 | Marinilabiliaceae - unknown genus |
|  |  | Hydrogenedentiales_31 | Hydrogenedensaceae - unknown genus |
|  |  | Erysipelotrichales_30 | Erysipelotrichaceae - Turicibacter |
|  |  | Coprothermobacterales_43 | Coprothermobacteraceae - Coprothermobacter |
|  |  | Clostridiales_26 | Peptostreptococcaceae - Intestinibacter |
| 3 | Grass | Clostridiales_10 | Lachnospiraceae - unknown genus |
|  |  | Methanofollis_64 | Methanomicrobiaceae - Methanofollis |
|  |  | Clostridiales_17 | Syntrophomonadaceae - Syntrophomonas |
|  |  | Clostridiales_42 | Peptococcaceae - Pelotomaculum |
|  |  | Clostridiales_77 | Clostridiaceae 1 - Clostridium sensu stricto 1 |
|  |  | Bacteroidales_67 | Rikenellaceae - DMER64 |
|  |  | Lactobacillales_87 | Enterococcaceae - Enterococcus |
|  |  | Lactobacillales_52 | Leuconostocaceae - Weissella |
|  |  | Bacteroidales_76 | M2PB4-65 termite group - unknown genus |
|  |  | Spirochaetales_79 | Spirochaetaceae - Treponema 2 |
|  |  | Clostridiales_80 | Clostridiaceae 1 - Clostridium sensu stricto 12 |
|  |  | Enterobacteriales_104 | Enterobacteriaceae - unknown genus |
| 4 | Fish | Methanosarcina_6 | Methanosarcinaceae - Methanosarcina |
|  |  | Clostridiales_2 | Clostridiaceae 1 - Clostridium sensu stricto 11 |
|  |  | Clostridiales_5 | Clostridiaceae 1 - Clostridium sensu stricto 13 |
| 5 | Fish and Sludge | Methanoculleus_18 | Methanomicrobiaceae - Methanoculleus |
|  |  | Clostridiales_15 | Syntrophomonadaceae - Syntrophomonas |
|  |  | Methanoculleus_50 | Methanomicrobiaceae - Methanoculleus |
|  |  | Methanobacterium_63 | Methanobacteriaceae - Methanobacterium |
|  |  | Methanobacterium_204 | Methanobacteriaceae - Methanobacterium |
